# O-GlcNAc modification of oncogenic transcription factor Sox2 promotes protein stability and regulates self-renewal in pancreatic cancer

**DOI:** 10.1101/345223

**Authors:** Nikita S Sharma, Vineet K Gupta, Patricia Dauer, Kousik Kesh, Roey Hadad, Bhuwan Giri, Anjali Chandra, Vikas Dudeja, Chad Slawson, Santanu Banerjee, Selwyn M Vickers, Ashok Saluja, Sulagna Banerjee

**Affiliations:** Department of Surgery, University of Miami, Miami, FL.; Department of Pharmacology, University of Minnesota, Minneapolis Minnesota.; University of Kansas Medical Center, Kansas City, KS.; Sylvester Comprehensive Cancer Center, Miami, FL.; Department of Psychology, Harvard University.; School of Medicine Dean’s Office, University of Alabama at Birmingham.

**Keywords:** OGT, O-GlcNAc, Sox2, Pancreatic cancer, metabolism, self-renewal

## Abstract

Pancreatic cancer is among the 3^rd^ leading cause of cancer related deaths in the United States along with a 5-year survival rate of 7%. The aggressive biology of the disease is responsible for such dismal outcome and is manifested by an increase in self-renewal capacity of the cancer cells, which leads to an increased rate of tumor-recurrence, contributing to poor prognosis. Transcription factor SOX2 maintains a critical balance between differentiation and “stemness” and is thus tightly regulated within a cell. In cancer, SOX2 is aberrantly “turned-on” leading to activation of self-renewal pathways in cancer. Regulation of Sox2 in cancer is poorly understood. In the current study, we show for the first time that in pancreatic cancer, Sox2 is modified by addition of O-GlcNAc moiety, catalyzed by OGT (O-GlcNAc Transferase) at S246. This activates Sox2 transcriptional activity by stabilizing the protein in the nucleus. A CRISPR-OGT knockout in pancreatic cancer cell line S2VP10 resulted in a delayed tumor initiation. We further showed that mutation of this site (S246A) prevents the modification of Sox2 and its downstream activity. Our study also demonstrated that targeting OGT *in vivo* with a small molecule inhibitor OSMI, results in decreased tumor burden, delayed tumor progression and a decreased expression of SOX2 in pancreatic cancer cells. Our study highlights for the first time that that the O-GlcNAc transferase dependent SOX2 glycosylation has a profound effect on the transcriptional activity of SOX2 and is instrumental in determining self-renewal in pancreatic cancer.

**Significance:** Our study highlights for the first time that that the O-GlcNAc transferase dependent SOX2 glycosylation determines self-renewal in pancreatic cancer which is responsible for tumor initiation.

## Introduction

Tumor initiating cells (TIC), a small population of cells within a tumor capable of turning on self-renewal properties, are thought to be responsible for tumor recurrence, metastasis and chemoresistance in a tumor. While there has been extensive research on this population of cells in pancreatic cancer and several studies have associated this population with poor prognosis, the molecular mechanisms of how these self-renewal genes are activated has remained an enigma. Self-renewal in any cell is regulated by a concerted action of the “Yamanaka factors” SOX2, OCT4, Myc and KLF4 (1). These transcription factors are robustly regulated by a number of interacting signaling pathways as well as cellular metabolism (2).Alteration in tumor metabolism and/or bioenergetics pathways in cancer have recently emerged as a new cancer hallmark (3). Cancer cells undergo drastic metabolic reprogramming that not only results in the classic Warburg Effect producing a large amount of lactate, but also in channeling a large amount of metabolites through biosynthetic pathways that support rapid growth and proliferation (4). Among these, the hexosamine biosynthesis pathway or HBP has emerged as a critical node that is instrumental in the regulation of a number of pro-oncogenic pathways (5-7). This pathway, often considered as a “nutrient-sensing” pathway, is a shunt pathway of glycolysis, uses fructose-6- and glutamine to produce glucosamine-6-phosphate. About 2-3% of the total cellular glucose is channelized via HBP to culminate in synthesis of the nucleic acid sugar UDP-GlcNAc. UDP-GlcNAc is one of the major substrates for both N-linked glycosylation as well as O-GlcNAcylation of proteins. Since the HBP is dependent on the availability of glucose, slight increases in glucose uptake results in significantly more O-GlcNAcylated modifications of many proteins (8-10).

Protein O-GlcNAcylation is mediated via the activity of the enzyme O-GlcNAc Transferase (OGT). A large number of proteins, particularly transcription factors are modified by OGT. This modification affects subcellular localization, protein-protein interaction or DNA binding ability of these proteins (11). Research from our group has shown that modification of Sp1 by OGT facilitates its nuclear translocation and thus regulates its transcriptional activity (5). A number of studies have been focused on how O-GlcNAc modification of pro-survival transcription factors may regulate tumor progression (11). Even though the O-GlcNAc modifications of transcription factors are well known, the regulation of self-renewal genes in cancer by O-GlcNAc modification is a relatively understudied area. Recent reports have shown that SOX2 and OCT4, the key transcription factors that regulate self-renewal in embryonic stem cells and induced pluripotent stem cells are modified by O-GlcNAc (12). In a recent study by Myers et al, O-GlcNAc modification of SOX2 regulates its interaction with PARP, which in turn led to differentiation (12). However, there is no report on how O-GlcNAc modification of SOX2 regulates the self-renewal properties in a tumor cell.

Hexosamine biosynthesis pathway, and particularly OGT expression is upregulated in pancreatic cancer (6,13,14). Studies from our laboratory have shown that O-GlcNAc modification of Sp1 regulates its nuclear translocation and thus its activity (5). We have also shown that O-GlcNAc modification of β-catenin in pancreatic cancer regulates the Wnt signaling pathway (15). In the current study, we have evaluated the role of O-GlcNAc modification on SOX2, the key transcription factor regulating self-renewal in cancer cells. Our results show that SOX2 is modified by O-GlcNAc transferase. O-GlcNAcylation of SOX2 stabilizes the protein in the nucleus and thus regulates its transcriptional activity. Further, mutating the Sox2 glycosylation site (S246A) prevents its O-GlcNAc modification and enhances its degradation. Inhibition of OGT in pancreatic cancer cells leads to a decrease in tumor initiation both *in vivo* and clonogenicity *in vitro*. This is the first report describing the O-GlcNAc modification of SOX2 in cancer cells and its role in regulation of self-renewal.

## MATERIALS AND METHODS

### Cell Culture

S2-VP10, and L3.6pL (a generous gift from Dr. Masato Yamamoto, University of Minnesota) cells were grown in RPMI 1640 - Gibco) and DMEM without NEAA containing 10% Fetal Bovine Serum (FBS) and 1% Penicillin/Streptomycin respectively. SU86.86 (ATCC) cell line was also cultured in RPMI 1640 (Gibco) with 10% FBS and 1% Penicillin/Streptomycin. Human pancreatic stellate cells (ATCC, Manassas, VA, USA) were cultured in stellate cell medium (ScienCell, 5301) with stellate cell growth supplements provided by the manufacturer (SteCGS, Cat. No. 5352, P/S, Cat. No. 0503) ScienCell, Carlsbad, CA, USA. All cell lines were routinely tested for mycoplasma contamination and STR profiles (ATCC).

### Transcriptome deep sequencing and analysis

Aliquots of RNA were derived from the qRT-PCR samples. 3 independent passages of OGTi and the respective sham shcells were analyzed. The RNA was quality tested using a Bioanalyzer 2100 (Agilent Technologies, CA, USA). cDNA was created by reverse transcription of oligo-dT purified polyadenylated RNA and fragmented, blunt-ended, and then ligated to barcoded adaptors. Then, the library was size selected, and the selection process was validated and quantified by capillary electrophoresis and qPCR, respectively. Samples were load on the HiSeq 2500 (Illumina Inc., CA, USA) to generate around 34 million paired-end 50bp reads for each sample.

Raw sequence data was processed through PartekFlow RNAseq pipeline (Partek Inc., St. Louis, MO) as follows- A pre-alignment QA/QC was performed and bases with Q>30 were retained for analysis. Bowtie2 was used to filter out non-human DNA, mtDNA and rDNA from the samples and STAR 2.5.3 aligner was used to map the high-quality reads to Hg19 human genome assembly. Aligned reads were quantified for differential abundance among samples using (a) Partek E/M annotation model and (b) Cufflinks algorithm using the Hg19- Ensemble transcripts release 75.

### Site Directed Mutagenesis

pDONR223_SOX2_WT obtained from Addgene (plasmid # 82233). SOX2 was modified at S246 to Alanine using QC Lightning Multi SITE DIRECTED MUTAGENESIS kit from Agilent according to manufacturer’s instruction. Clones were confirmed by sequencing (16).

### Immunofluorescence

Tissues were de paraffanized by heating it at 56°C overnight and then hydrated by treating it with xylene (15 mins, two times),100% ethanol,90% ethanol, 70% ethanol (2 times) 5 mins each. The slides were then steamed with a pH 6 reveal decloaker (BIOCARE Medical, Concord, CA, USA) for antigen retrieval, blocked in Dako serum blocker (Agilent technologies, Santa Clara, CA, USA). Primary antibody was added overnight. Slides were washed 3X in PBS, secondary antibodies (Alexaflour) were diluted in SNIPER (BIOCARE Medical) and slides were stained for 30 mins at room temperature. Slides were washed again 3X in PBS and mounted using Prolong Gold antifade with DAPI (Molecular Probes, Thermofischer Scientific, Weston, FL,USA). Slides were dried overnight and imaged by fluorescent microscope. SOX2 antibody was purchased from SIGMAALDRICH, St Louis, MO, USA (SAB2701974), OCT 4 (D7O5Z) and NANOG (1E6C4) antibody was purchased from CELL SIGNALLING TECHNOLOGY, Danvers, MA, USA. Antibody against OGT and O GlcNAc were purchased from SIGMA-ALDRICH (SAB2101676) and Millipore, Burlington, MA, USA (Clone CTD 110.6), respectively. Images were acquired at 20X and 60X

### Generation of OGT-Knockout Cell line

Human OGT gene knockout kit was obtained from Origene, Rockville, MD, USA (KN206822). S2-VP10 cells were plated at a density of 80,000 in a 6 -well plate. One day after plating the cells were transfected with gRNA1 and gRNA2 along with donor vector. The cells were then cultured according to the manufacturer’s protocols. After 8 passages puromycin pressure at a concentration of 0.50μg/ml was applied. The colonies were allowed to grow and the cells were then single-cell sorted by FACS in a 96-well plate. The select clones were allowed to grow and then characterized by mRNA expression and western blot.

### Proximal Ligation Assay

S2VP10 and L3.6pL cells were plated at a density of 50,000 cell per well in a chamber slide. Cells were treated with OSMI for 24 hours. After 24 hour’s media was removed, the cells were washed 3 times with PBS and fixed with 2% paraformaldehyde. After fixing the cells, they were stained according to the manufacturer’s protocol (Duo link, DU 092101, SIGMA) and the images were acquired at 20X magnification.

### *In vivo* Studies

Female athymic nude mice between the ages of 4-6 weeks were used for *in vivo* experiments and were purchased from the Jackson laboratory, Bar Harbour, ME, USA. For subcutaneous experiments 500,000 cells were implanted in the right flank of the mice. Corning^®^ Matrigel^®^ Growth Factor Reduced (GFR) Basement Membrane Matrix, purchased from Corning, inc, Corning NY, USA and 1X PBS at a ratio of 1:1 were used as a suspension medium for the cells. Tumors were allowed to reach a size of 100mm^3^ and then treatment was started. OSMI (SML1621 SIGMA) was given at a dose of 10mg/kg/day in DMSO for 3 weeks. At the end of 3 week’s mice were sacrificed and tissues were collected.

For tumor initiation studies, OGT knockout (OGTi-S2VP10) cells were injected at a limiting dilution from 500,000 cells per mice to 500 cells in the right flank of female athymic nude mice. The mice were monitored daily and the appearance of tumor was noted. All the experiments were performed according to the IACUC protocols as approved by the University of Miami.

### RNA Isolation and cDNA synthesis

RNA was isolated using the Trizol method of isolation and cDNA was made using the High Capacity cDNA Reverse Transcription Kit. 2 µgm of RNA was used per reaction. RT-qPCR was performed in the Roche Light Cycle 480.Primers (SOX2 PPH02471A, OCT4 PPH66786A, NANOG PPH17032E-200, OGT PPH19166A, OGA PPH01061A) were purchased from Qiagen, Valencia, CA, USA

### Western Blotting

Cell lysates were prepared using RIPA lysis buffer with protease and phosphatase inhibitors. 4-20% SDS page gel were used for running the proteins.5% BSA in TBST was used as a blocking agent. SOX2 antibody (SAB2701974) was purchased from SIGMA-ALDRICH and was used at a dilution of 1:1000. Secondary anti rabbit was used at a dilution of 1:5000.

### Sox2 Dual Luciferase Reporter Assay

S2VP10 and L3.6pL cells were plated at a density of 0.8×10^6^ cells per well in a 24 well plate. Cells were transfected with the SOX2 reporter plasmid the next day using Attractene as a transfection reagent. The day after the transfection with the reporter plasmid the cells were treated with OSMI and/ or MG132 at a dose of 50µM and 1µM respectively for another 24 hours respectively. 24 hours after treatment the cells were lysed and the luminescence was analyzed using a spectrophotometer.

### Sox2 ELISA

For estimating the amount of SOX2 protein in our cell lysates, we performed a Sandwich ELISA using the human SOX2 ELISA kit from My Biosource, San Diego, CA, USA (MBS2513016). Cells were plated in a 10 cm dish and allowed to attach and grow for 24 hours. After 24 hours of plating cells were treated with OSMI and MG1332 alone or in combination at a dose of 50 µM and 1µM for 24 hours respectively. After 24 hours of treatment cells were lysed using RIPA lysis buffer and the protein was estimated using BCA protein estimation method. Equal amount of protein was loaded for the assay and the assay was performed according to the manufacturer’s protocol.

### Ethics Statement

All procedures were performed according to protocols approved by the University of Miami Institutional Animal Care and Use Committee (IACUC).

### Statistical Analysis

Values are expressed as the mean +/- SEM. All *in vitro* experiments were performed at least three times. The significance between any two samples was analyzed by t-test, values of p<0.05 were considered statistically significant.

## RESULTS

### OGT expression correlates with the aggressiveness of the disease

OGT expression and O-GlcNAc modifications are elevated in pancreatic cancer (17). Previously published results from our laboratory confirm this (5). To further investigate if this phenomenon is so in patient tumor tissues, we performed immunohistochemistry on a de-identified human pancreatic tumor tissue microarray. Our results showed that the expression of OGT increases with the aggressiveness of the disease. OGT expression was significantly higher in the T3N0M0 stage of the pancreatic tumor compared to the adjacent normal or the T2N0M0 stage (Figure 1A, B). To evaluate whether O-GlcNAcylation of proteins is increased as the pancreatic tumor progresses, we first compared the expression of OGT in the pancreas of 1 month old Kras^G12D^, TP53 ^R172H^ Pdx-^Cre^ (KPC) mice and 6-8-month-old KPC mice with a fully developed tumor. Our results showed that expression of OGT was significantly less in the tissue of 1-month-old mice (lacking tumor) compared to that in a 6-8month-old KPC mice (that has tumor) (Figure 1C). The TNM staging of the TMA is shown in Supplementary Figure 1A, B. To study whether this was also demonstrated in the patient databases, we performed an analysis of the OGT expression and correlated it with patient survival in the TCGA database at www.cbioportal.org. Our results showed a negative correlation between the expression of OGT and patient survival and advanced histologic grade (Supplementary Figure 1C, D).

**Figure 1.**
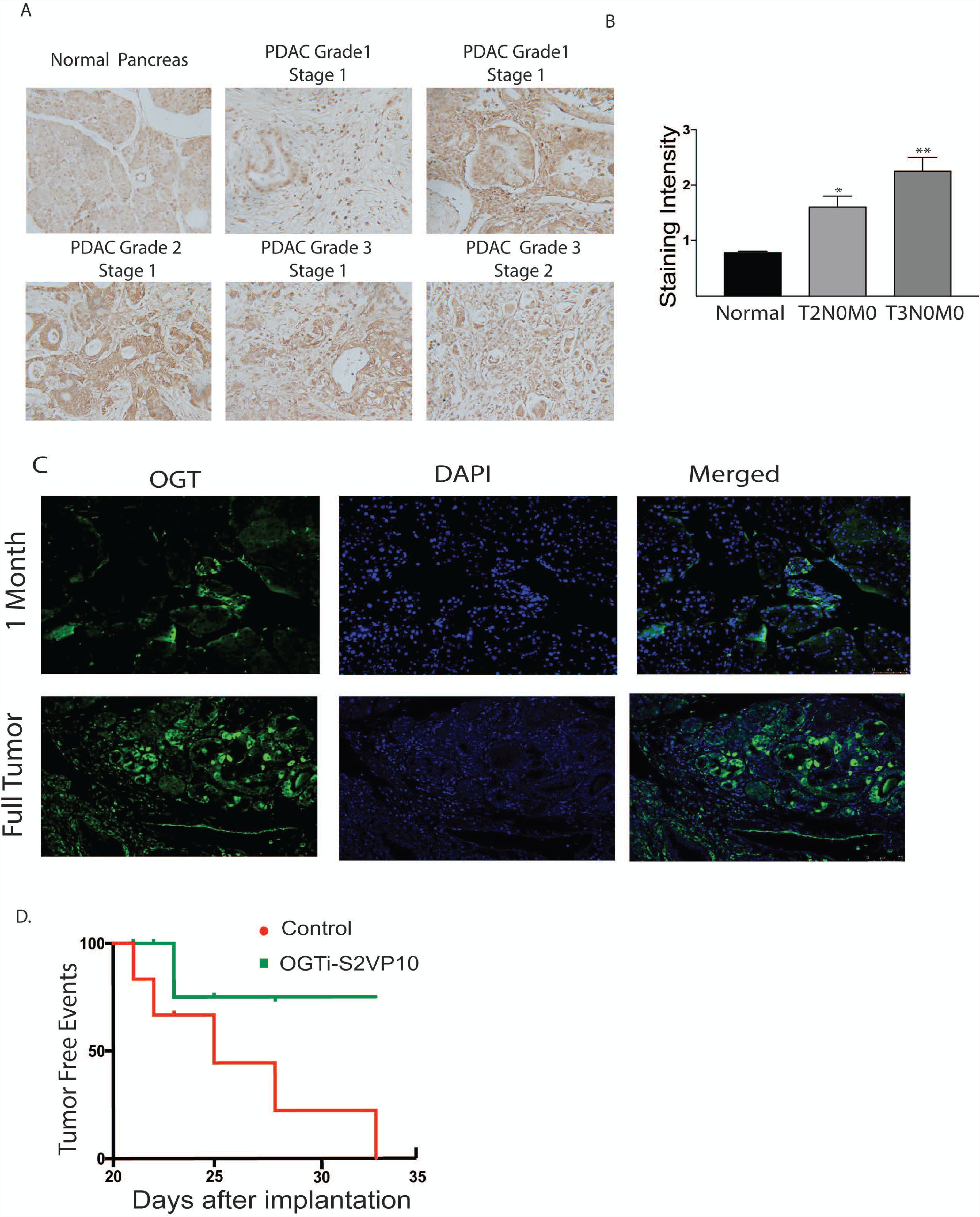
OGT expression in pancreatic cancer. Tumor tissue microarray from patient samples showed that OGT expression coincided with the aggressive stages of the disease (A). The quantitation of the histology confirmed this observation (B). In Kras^G12D^TP53^R172H^ Pdx-cre or KPC model for pancreatic cancer, OGT expression increased as the tumor progressed. Representative immunofluorescence from 1 month old mice and mice with full tumor (C). CRISPR knockout of OGT in pancreatic cancer cell line S2VP10 showed delayed tumor initiation compared to scrambled control (D).

Our data corroborated with previous findings that OGT and OGA are overexpressed in pancreatic cancer cell line (Supplementary Figure 1 E, F, G) (7). Since S2VP10 and L3.6 PL cell lines had the highest expression of OGT among those tested, we used these in the current study. Tumor aggressiveness is typically characterized by their ability to recur after resection or therapy. This phenomenon is characterized by the “turning-on” of self-renewal genes like SOX2, OCT4 and Nanog(1). To study the effect of OGT down-regulation on tumor recurrence, we used an OGT-CRISPR plasmid that stably knocked-out the expression of OGT in highly aggressive S2VP10 pancreatic cancer cells (OGTi). These cells showed decreased total O-GlcNAc level as well as decreased expression of OGT (Supplementary Figure 2A, B). When implanted in limiting dilution (500 cells) in athymic nude mice, they showed a distinct delay in tumor initiation suggesting a definite role in tumor recurrence (Figure 1D). This indicated that OGT was instrumental in tumor initiation of pancreatic cancer cells.

### Transcriptomic analysis of OGT knockout cell line

O-GlcNAc transferase or OGT is responsible for modification of a number of critical pathways that actively affect oncogenic signaling in cancer. To study the major pathways that are downregulated upon inhibition OGT, we performed a RNA-seq analysis on OGTi cells. Raw sequence data was processed through PartekFlow RNAseq pipeline (Partek Inc., St. Louis, MO) as follows- A pre-alignment QA/QC was performed and bases with Q>30 were retained for analysis. Bowtie2 was used to filter out non-human DNA, mtDNA and rDNA from the samples and STAR 2.5.3 aligner was used to map the high-quality reads to Hg19 human genome assembly. Aligned reads were quantified for differential abundance among samples using (a) Partek E/M annotation model and (b) Cufflinks algorithm using the Hg19- Ensemble transcripts release 75. Out of 13,401 identified transcripts, 43 were found to be significantly different among the two study groups. (Figure 2A, D). The Gene Ontology showed that the majority of the deregulated genes belonged to the category of embryogenesis, organogenesis and differentiation, cell survival and regulation of gene transcription (Figure 2B). To further analyze the pathways that are deregulated upon inhibition of OGT, we conducted pathway enrichment analysis and found that most significantly deregulated genes belonged to pathways were those involved in development and differentiation (Figure 2C, D). Among these, the human embryonic stem cell and pluripotency network was observed to be present. This network encompassed the downstream signaling stemming from PDGF-PDGFR and Wnt and culminated in regulation of genes that were involved in self-renewal pathways (Figure 2E).

**Figure 2.**
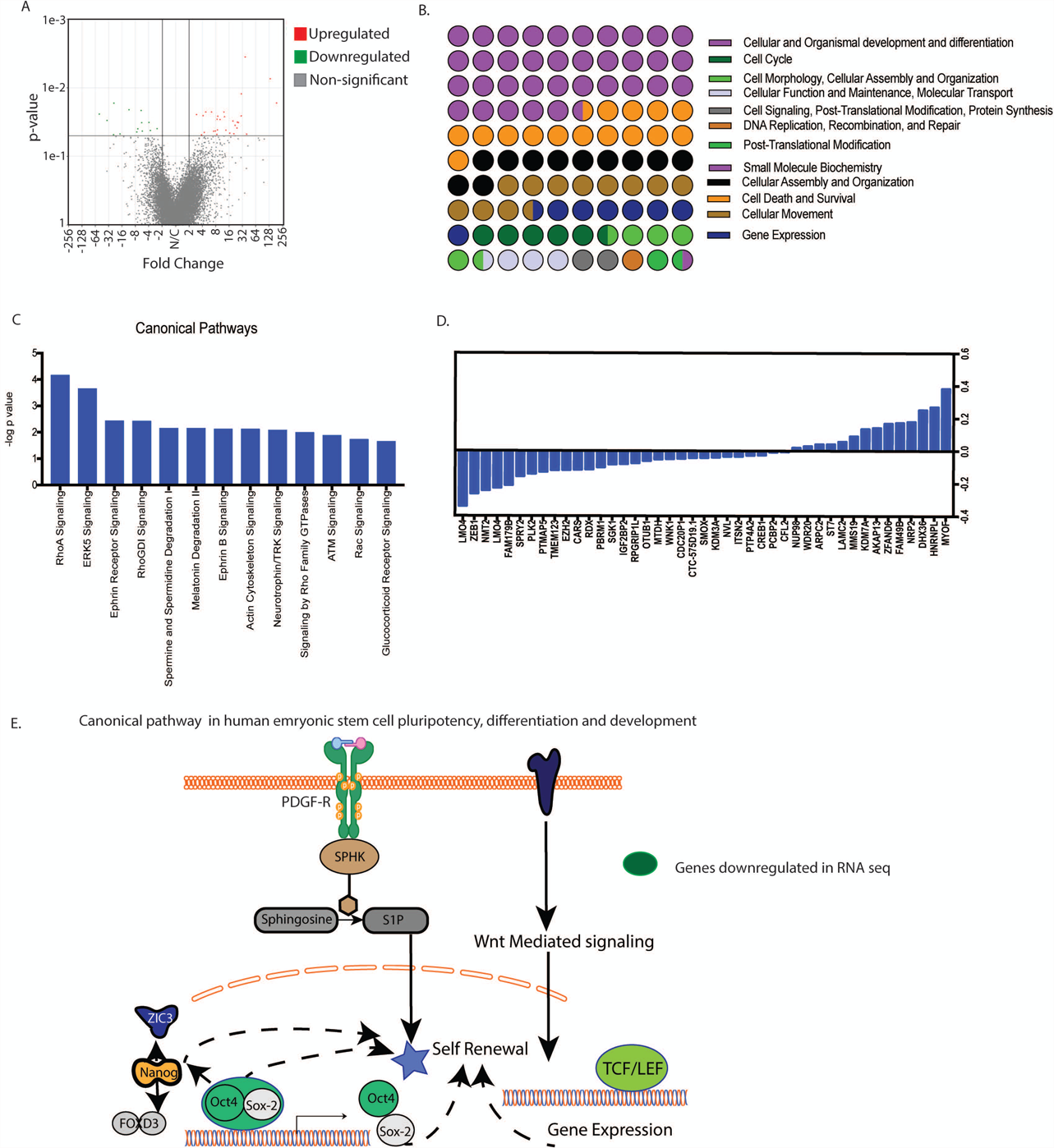
Transcriptomics analysis of OGTi cells: Volcano plot showing significantly deregulated genes (A) GO of the biological processes that are altered as a result of OGT inhibition (B). Canonical pathways altered by OGT inhibition as analyzed by the IPA software (C). List of 43 deregulated genes upon OGT inhibition (D). Schematic of embryonic stem cell pathways that are deregulated upon OGT inhibition (E),

### Down-regulation of OGT leads to a decrease in self-renewal in pancreatic cancer

Since OGT inhibition resulted in a delay in tumor initiation, we next studied the effect of this inhibition on self-renewal genes namely, SOX2, OCT4 and NANOG. Our results showed that OGT silencing (using siRNA) did not change the expression of core self-renewal transcription factors SOX2 or OCT 4 in S2VP10 cells while modestly changing its expression in L3.6pL cells (Supplementary Figure 2C, D). However, a methyl-cellulose based colony forming unit assay on S2-VP10 and L3.6pL cells showed that treatment with OSMI (a small molecule inhibitor of OGT) decreased the colonies in S2VP10 and L3.6pL cells (Figure 3 A,B) compared to untreated cells. Since small molecule inhibitors are notorious for their off-target effects, we next performed this assay using the OGTi-S2VP10 cells. The colony forming ability of these cells was also significantly decreased (Figure 3B). Since colony-forming ability is a measure of the cells self-renewal potential, we next performed a luciferase based transcriptional activity assay on the transcription factors regulating self-renewal. Pancreatic cancer cells S2VP10, were treated with small molecule inhibitor OSMI and used for this luciferase based stem cell reporter array. Our results showed that among the 10 transcription factors involved in self-renewal SOX2 activity was the most significantly down-regulated (Supplementary Figure 3). We validated this using a Luciferase based transcriptional activity assay for SOX2, OCT4 and Nanog using OSMI (Figure 3C) and siOGT on S2VP10 cells (Figure 3D). SOX2 transcriptional activity assay was also performed on L3.6PL cells using siOGT (Figure 3D) and on our OGTi-S2VP10 cells (Figure 3E). Our results showed that SOX2 activity was affected significantly in cells in which OGT was inhibited either by OSMI or by siRNA compared to the control. Our results thus indicated that OGT is required for SOX2 transcriptional activity.

**Figure 3.**
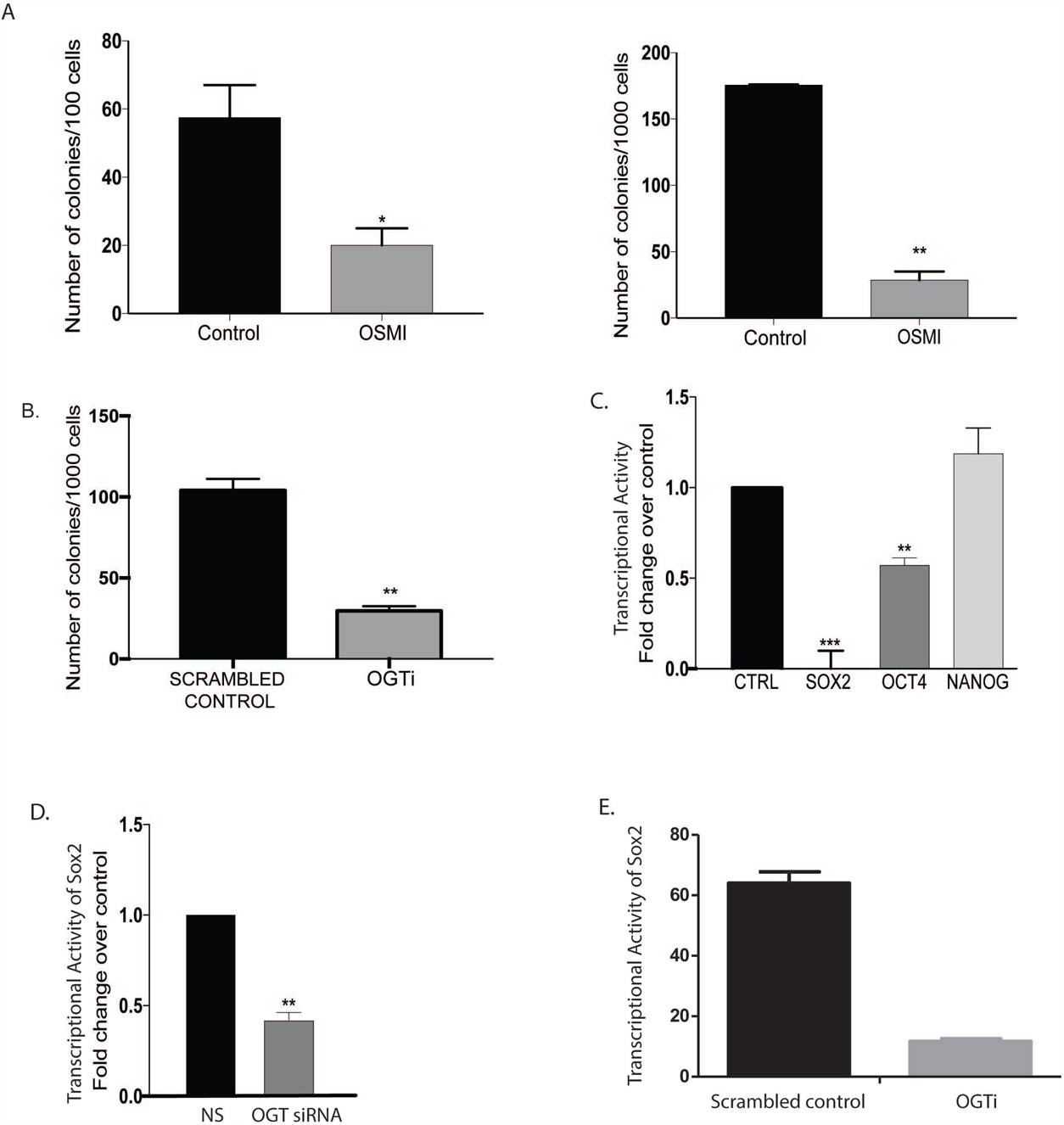
Downregulation of OGT expression leads to a decrease in self-renewal in pancreatic cancer: In S2VP10 (A) and L3.6pL cells (B) inhibition of OGT by small molecule inhibitor OSMI decreased the colony forming units compared to untreated indicating that OGT was instrumental in determining clonogenicity. CRISPR knockout of OGT in S2VP10 cells (OGTi) also showed decreased colony formation (C) compared to scrambled control. Transcriptional activity of Sox2 was decreased upon inhibition of OGT with an inhibitor, OSMI (C) and with siRNA (D). OGTi cells showed a similar decrease in the transcriptional activity of SOX2.

### Transcriptional regulation of self-renewal in pancreatic cancer is dependent on O-GlcNAc modification of Sox2

Since our results indicated that OGT inhibition might affect self-renewal properties of pancreatic cancer cells via inhibition of SOX2 activity, we next studied the glycosylation of SOX2. SOX2 is modified by OGT in embryonic stem cells. This modification of SOX2 is known to play a role in its protein-protein interaction in those cells (12). To determine if SOX2 is modified by O-GlcNAc in pancreatic cancer cells, we performed an immunoprecipitation with O-GlcNAc antibody and probed it with anti-SOX2 antibody. SOX2 was seen to be glycosylated by O-GlcNAc in this experiment (Figure 4A). To further confirm this, we performed a Proximal Ligation Assay (PLA) on S2-VP10 cells. Our results showed that SOX2 is indeed modified by O-GlcNAc and that inhibition of OGT by OSMI or siRNA caused a decrease in its glycosylation (Figure 4B). As seen with OSMI, immunofluorescence with SOX2 and O-GlcNAc in the stable KO of OGT in OGTi-S2VP10 also showed decreased glycosylation on SOX2 protein (Figure 4C).

**Figure 4.**
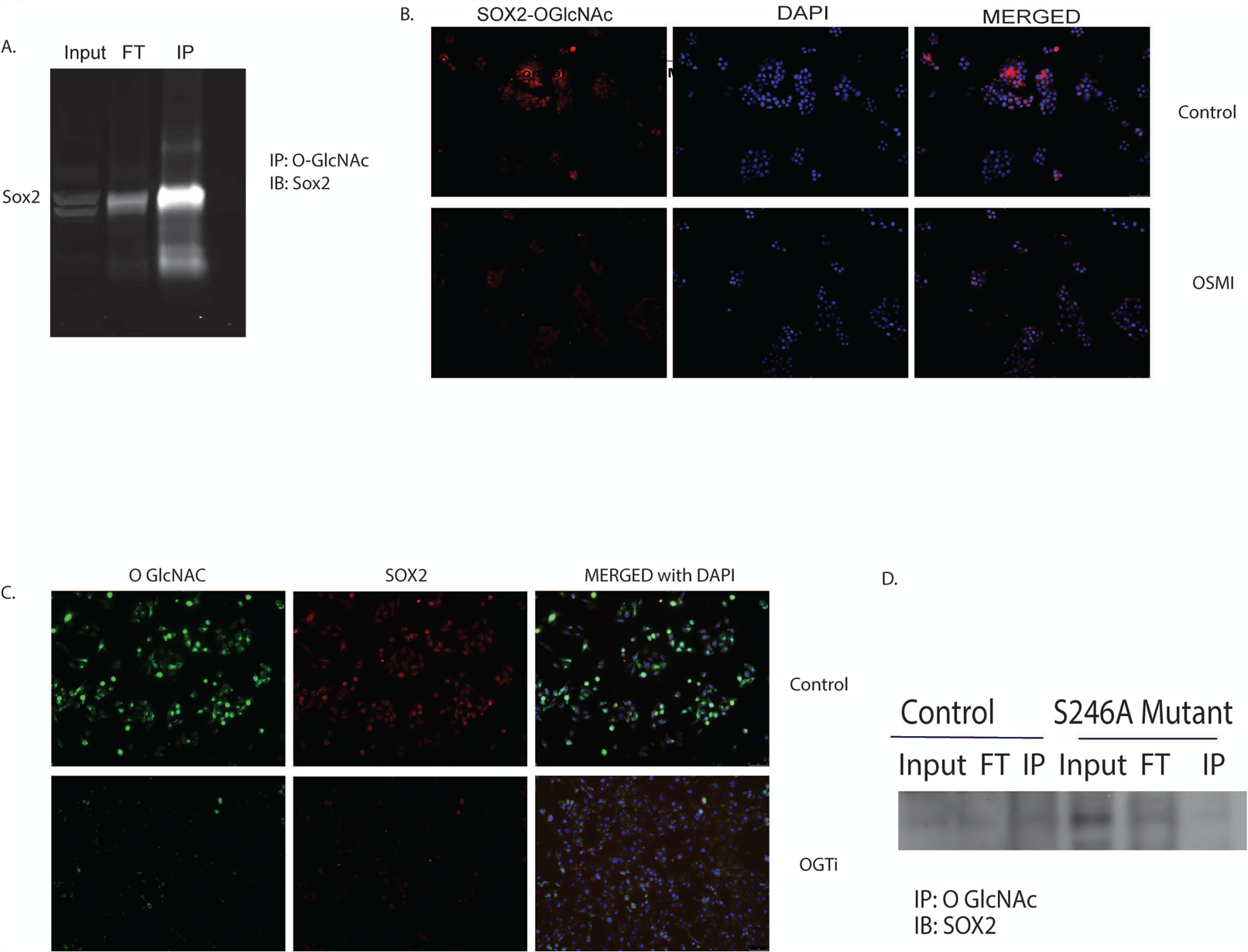
Transcriptional regulation of self-renewal in pancreatic cancer is dependent on O-GlcNAc modification of Sox2. Immunoprecipitation with anti-O-GlcNAc Ab followed by western blotting with Anti-Sox2 Ab showed that Sox2 was modified by addition of an O-GlcNAc moiety (A). Proximal ligation assay with Anti-O-GlcNAc ab and Anti-Sox2 Ab confirmed this modification. Upon treatment with 50uM OSMI, the modification was lost as seen by loss of fluorescence (B). In OGTi cells, there was decreased Sox2 expression compared to control as visualized by immunofluorescence (C). Immunoprecipitation of S2VP20 cells transfected with S246A mutated plasmid leads to a decrease in the interaction between Sox2 and O GlcNAc (D),

To study if the modification of Sox2 is disrupted by mutating the site we next performed a site directed mutagenesis of Ser 246 to Ala in Sox2 overexpressing plasmids obtained from Addgene. The plasmid was then transiently transfected to S2-VP10 cells for 24 hours. We observed that S246A mutation in Sox2 gene resulted in loss of O-GlcNAc modification of SOX2 as observed via immunoprecipitation (Figure 4D), confirming that Ser246 in SOX2 indeed gets modified by O GlcNAcylation.

### O-GlcNAc modification of SOX2 is required for its stability in pancreatic cancer cells

O-GlcNAc modification of a protein can affect its activity in a number of ways. In embryonic cells, modification of SOX2 facilitates its interaction with PARP and promotes differentiation (12). Our results (in Figure 3) showed that upon inhibition of OGT, SOX2 transcriptional activity was inhibited. This could be because OGT regulates DNA binding ability of SOX2. To determine this, we studied if inhibition of OGT affected the DNA binding ability of SOX2. Our results indicated that the DNA binding ability of SOX2 is inhibited in OGTi cells (Figure 5A) as well as in L3.6PL cells when treated with OSMI (Figure 5B). To study if this inhibition was owing to decreased SOX2 protein level in cells, we next studied the protein expression of SOX2 upon OGT inhibition. Our results showed that SOX2 protein was indeed decreased upon OGT inhibition by OSMI or genetic inhibition of OGT by CRISPR-OGTi (Figure 5 C, D). Since our observation (in Supplementary Figure 2C) indicated that SOX2 mRNA remained unaltered, we concluded that inhibition of OGT was affecting the stability of SOX2 protein. It is well known that SOX2 acts as a molecular rheostat for a cancer cell and its expression needs to be tightly regulated. Thus, the degradation machinery for SOX2 needs to be fairly robust in a cell (18). To further validate this result, we treated S2VP10 and L3.6pL cells with OSMI alone and OSMI with MG-132, a proteasome inhibitor. We observed a significant rescue in the protein levels of SOX2 when the cells were treated with the proteasome inhibitor (Figure 5E) thus demonstrating that O-GlcNAcylation of SOX2 is important for its stability. Further, treatment with MG132 in the presence of OSMI also showed a rescue in the transcriptional activity of SOX2 (Figure 5F). Further, our studies showed that mutation of the OGlcNAc modification site (S246A) resulted in a profound decrease in the SOX2 protein levels indicating that loss of O-GlcNAc modification of Sox2 led to decreased protein (Figure 5 G) and reduced Sox2 DNA binding activity (Figure 5H). We also observed that upon inhibition of OGT (by treatment with OSMI), there was an inhibition of nuclear translocation which was reverted upon treating with O-GlcNAcase inhibitor Thiamet G (Figure 5I). Additionally, quantitation of nuclear and cytosolic Sox2 in OGTi cells showed the same results (Figure 5J).

**Figure 5.**
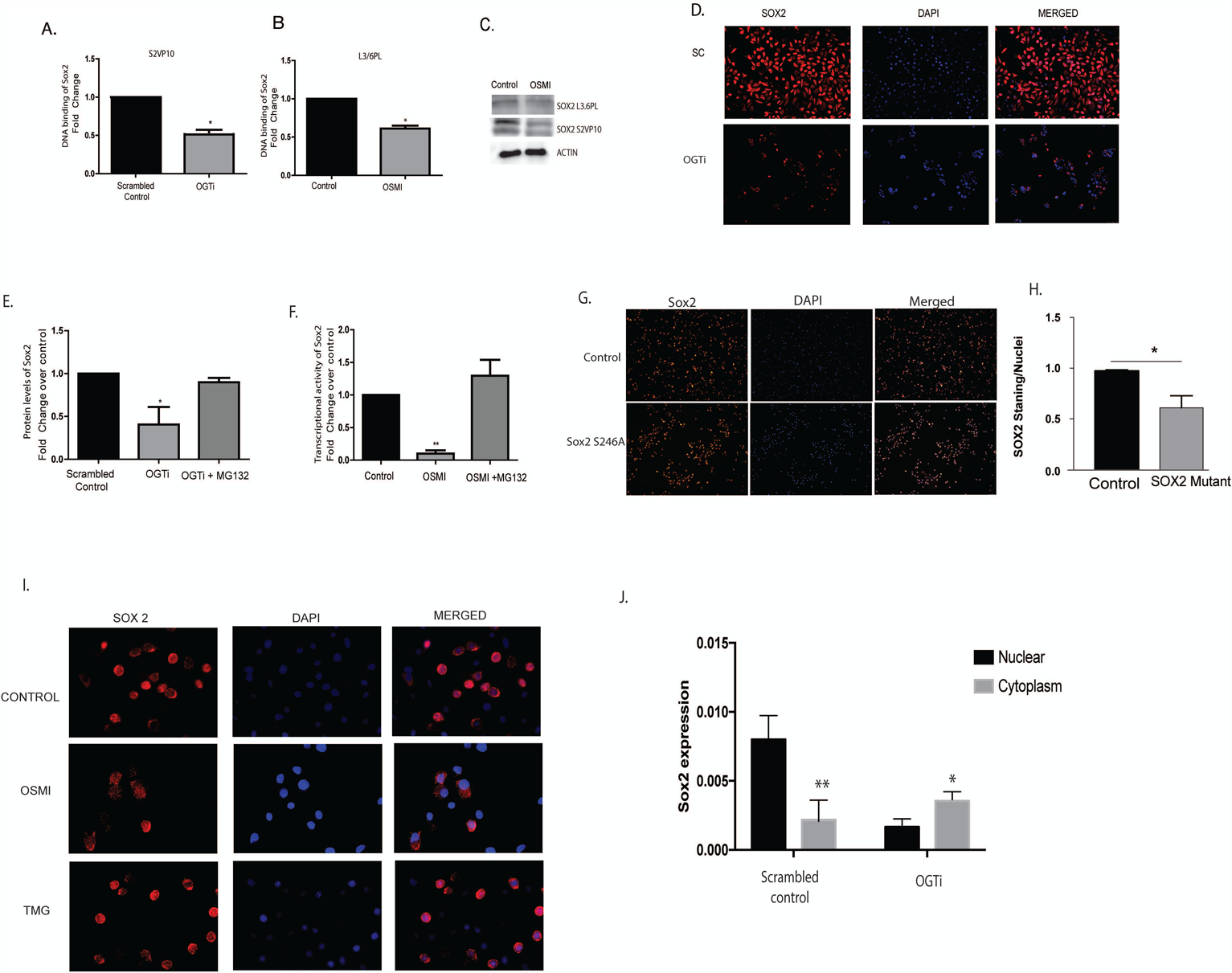
O-GlcNAc modification of SOX2 is required for its stability in pancreatic cancer cells. Inhibition of OGT in the CRISPR-OGTi cells resulted in decreased DNA binding by Sox2 proteins (A). Similarly, in L3.6 PL cells, treatment with OSMI decreased Sox2 DNA binding (B). Treatment with OSMI 50uM also decreased Sox2 protein level as seen by western blot (C) and immunofluorescence (D). Upon treatment with a proteosomal inhibitor MG132 in the presence of OSMI, Sox2 protein levels were restored in the pancreatic cancer cells (E) as estimated by the SOX2 protein ELISA. Treatment with MG132 in the presence of OSMI also rescued Sox2 transcriptional activity as seen by a luciferase based reporter assay (F). Transient transfection of S2VP10 cells with a SOX2 S246A mutant plasmid lead to a decrease in the protein levels of SOX2 (G, H). Treatment with OSMI (50uM), resulted in inhibition of nuclear translocation of Sox2 while inhibition of OGA (the glycosidase that mediates removal of O-GlcNAc from a protein) using Thiamet-G (TMG), resulted in increased nuclear translocation compared to untreated cells (I, J).

### Inhibition of OGT *in vivo* decreases tumor burden

To evaluate the efficacy of OGT downregulation *in vivo*, we implanted S2VP10 cells subcutaneously in athymic nude mice. OSMI was administered at a dose of 10mg/kg/day for 30 days. Consistent with our *in vitro* data treatment with OSMI caused a delay in tumor progression (Figure 6A). Further, a significant decrease in tumor volume and weight was observed upon inhibition of OGT (Figure 6 B, C). The tumors showed less O-GlcNAc (Figure 6D) and decreased cellularity in the H&E staining (Figure 6E). Further, as expected, there was less O-GlcNAc modification on SOX2 (Figure 6F). Upon analysis, the tumors showed a decrease in the OCT4 and NANOG expression when treated with OSMI (Supplementary Figure 4).

**Figure 6.**
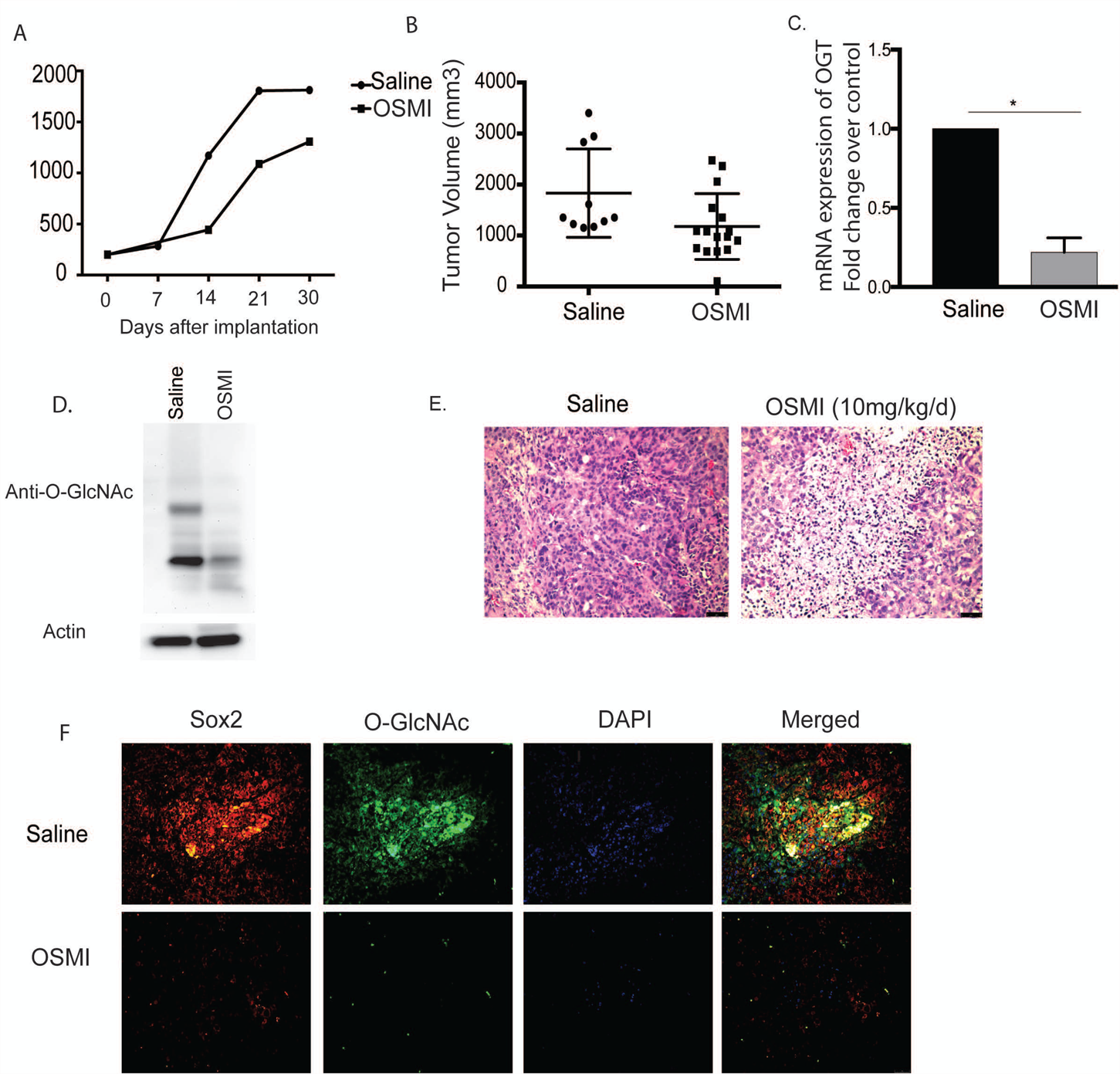
Inhibition of OGT *in vivo* decreases tumor burden. Treatment with OGT inhibitor OSMI delayed tumor progression (A) and decreased tumor volume (B). Treatment with OSMI decreased mRNA expression of OGT in the tumors (C) and total O-GlcNAcylation (D). Tumors treated with OSMI also showed decreased cellularity as seen by H&E stains in immunohistochemistry (E). Sox2 protein levels and O-GlcNAcyltion was also decreased upon treatment with OSMI (F).

## Discussion

O-GlcNAc transferase or OGT is an enzyme that transfers a single N-Acetyl glucosamine to serine or threonine residues of many proteins. This modification can “flag” the protein for phosphorylation (19), determine its interaction with other proteins or regulate its nuclear translocation (11,20). The role of O-GlcNAc modification on key oncogenic transcription factors has gained importance over the last decade (11,21). A number of proteins like Myc, Sp1 and NF-kB which are instrumental in promoting proliferation in cancer, are known to be modified by O-GlcNAc (5,7). Previous results from our laboratory have shown that O-GlcNAc modification of Sp1 regulates its nuclear translocation and thus its activity (5). Similarly, recent studies have shown that O-GlcNAc modification of the YAP component of the Hippo Pathway disrupts its interaction with LATS1 (an upstream kinase) thereby preventing its phosphorylation and up-regulating its transcriptional activity (22). In gastric cancer, O-GlcNAc modification stabilizes FOXM1 protein by decreasing its phosphorylation by GSK-3β (23).

The role of O-GlcNAc modifications in regulating stemness and self-renewal is gaining importance in cancer research (24,25). A recent study in embryonic stem cells has shown that Sox2, a key transcription factor for self-renewal is modified by O-GlcNAc, which in turn facilitates its interaction with PARP to regulate differentiation (12). However, there are very few studies on whether this modification is responsible for initiation and regulation of self-renewal in cancer. Our current study shows that inhibition of OGT by a CRISPR results in a delay in tumor formation in animals (Figure 1). Tumor initiation is regulated by SOX2, OCT4 and NANOG transcription factors. These transcription factors act in a concerted manner to regulate genes that control “stemness” and differentiation and thus are responsible for recurrence of tumor. Our study shows that in pancreatic cancer, where there is a high incidence of tumor recurrence, SOX2 was getting modified by O-GlcNAc (Figure 4).

O-GlcNAcylation of SOX2 regulates its transcriptional activity. When OGT is inhibited either by siRNA, or the small molecule inhibitor OSMI, there is a distinct decrease in colony formation units (Figure 3). Further, tumors treated with OSMI *in vivo* showed a clear decrease in the expression of SOX2 (Figure 6). Our results also showed that O-GlcNAc modification was decreasing SOX2 activity by affecting its stability and preventing its nuclear localization (Figure 5). This was an interesting observation as it had been reported earlier that in embryonic stem cells, the acetylation Sox2 regulates its nuclear translocation (26,27). However, our studies clearly indicated that upon inhibition of OGT, there is an inhibition of nuclear translocation when O-GlcNAc modification of SOX2 is abrogated. Conversely, treatment of pancreatic cancer cells with Thiamet-G, an inhibitor of O-GlcNAse (an enzyme that facilitates removal of glycosylation) reverted this and promoted nuclear translocation. This indicates that SOX2 glycosylation regulates its nuclear translocation (Figure 6B). It is possible that the regulation of sub-cellular localization of SOX2, is not directly regulated by OGT, but is via other routes (for example, activating the acetylation of Sox2). However, studying that is beyond the scope of the current manuscript.

The stability of SOX2 has been implicated to play a role in its function. Sumoylation of SOX2 at Lysine 247 is instrumental in inhibition of DNA binding by SOX2 (28). Our results showed that OGlcNAc modification of SOX2 also regulates its stability (Figure 5). In a number of other cancers, O-GlcNAc modification is known to regulate protein stability. In prostate cancer, O-GlcNAc modification of Bm1-1 modulates its stability and promotes oncogenic function (29,30). However, its role in stability of SOX2 protein in pancreatic cancer cells has not been studied before. Our study thus show for the first time that O-GlcNAc modification in pancreatic cancer may be instrumental in regulating its self-renewal properties by maintaining the stability of Sox2 and regulating its nuclear translocation.

## Conclusion

Self-renewal in a tumor is considered to be one of the hallmarks of the cancer stem cells or tumor initiating cells that are responsible for tumor recurrence, chemoresistance and metastasis. Understanding the molecular switch that can potentially turn-on the self-renewal pathway (by activating Sox2), is thus a significant area of research. In this context, our study highlights that the O-GlcNAc transferase dependent SOX2 glycosylation has a profound effect on the transcriptional activity of SOX2 as this modification regulates its nuclear translocation as well as protein stability. Understanding this mechanism of “turning-on” Sox2 by its glycosylation is thus expected to pave the way for development of novel therapy that can not only decrease tumor progression but also eradicate the “roots” of cancer or the “cancer stem cells”.

## Figure Legends

**Supplementary Figure 1.**
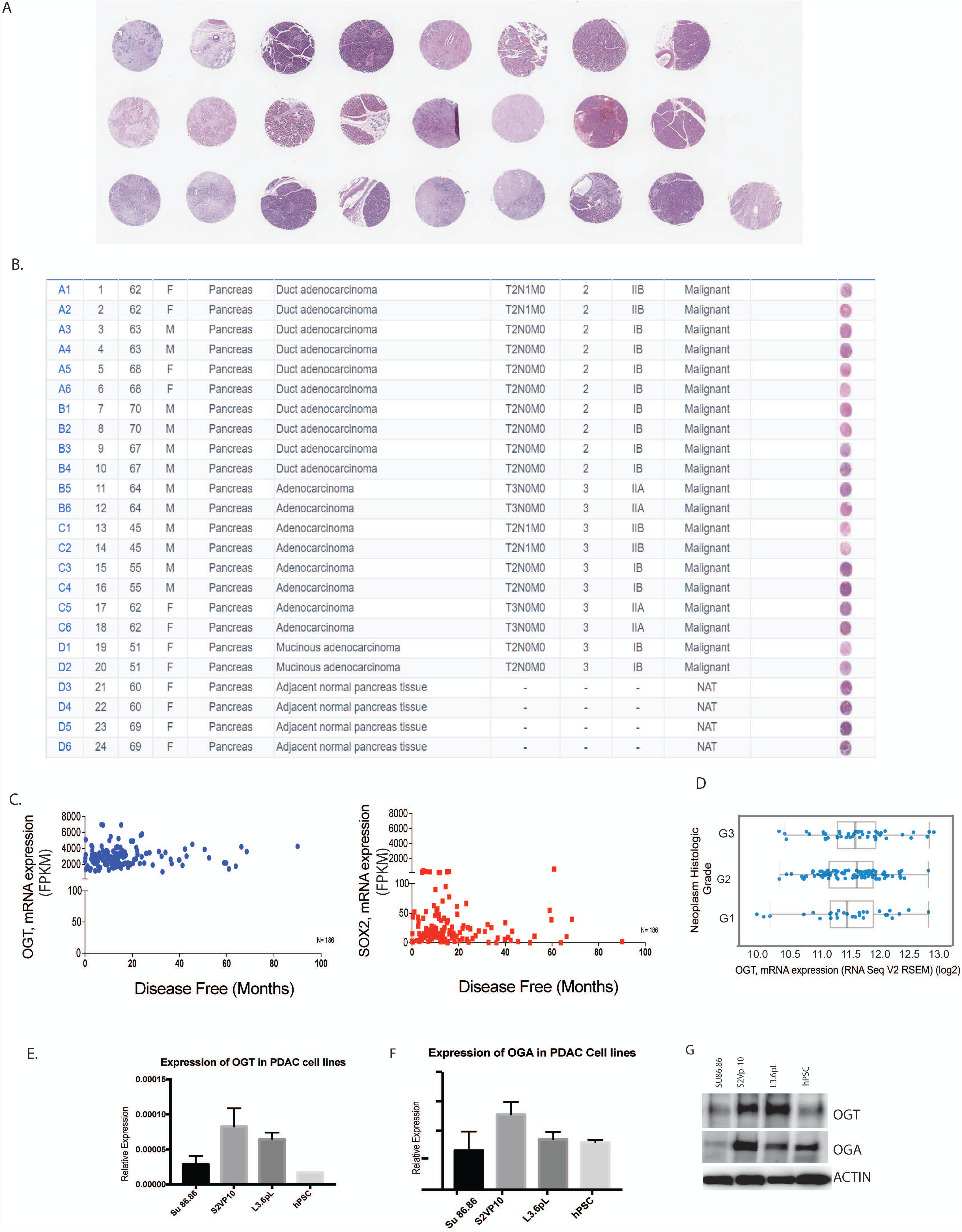
Images of tumor tissue microarray (A) with TNM staging (B). Analysis of samples in TCGA at www.cbioportal.org showed that high OGT and Sox2 expression correlated with less disease free survival (C). Further, histologic grade showed increased OGT expreesion correlated with advanced histologic grade (D) RNA expression of OGT (E), OGA(F) in pancreatic cancer cell lines; Protein expression of OGT and OGA in pancreatic cancer cell lines (G).

**Supplementary Figure 2.**
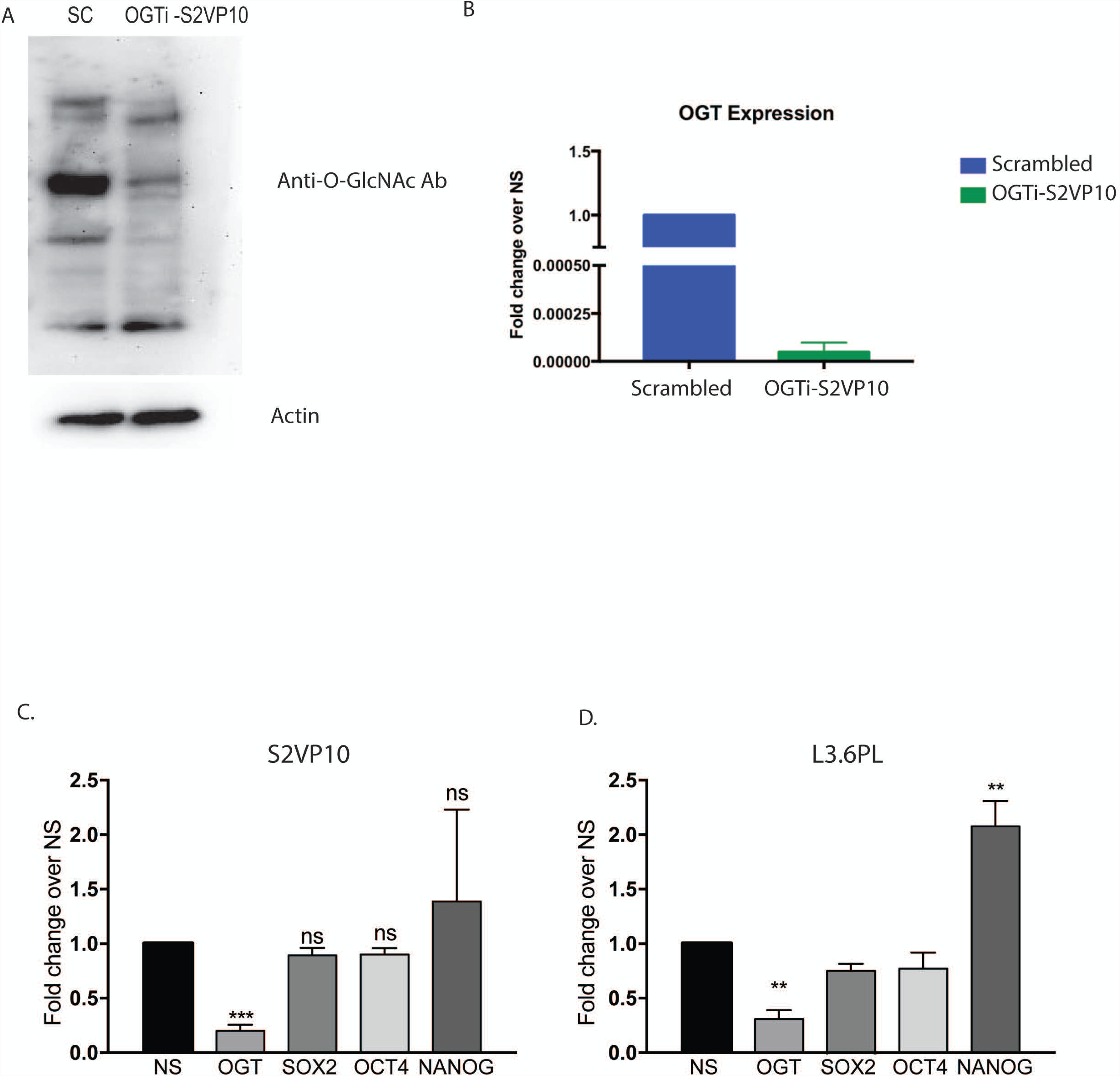
CRISPR-OGT knockout (OGTi) shows decreased O-GIcNAcation (A) as well as decreased RNA expression of OGT (B). Downregulation of OGT in pancreatic cancer cell S2VP10 (C) or L3.6PL (D) did not affect mRNA expression of self renewal genes.

**Supplementary Figure 3.**
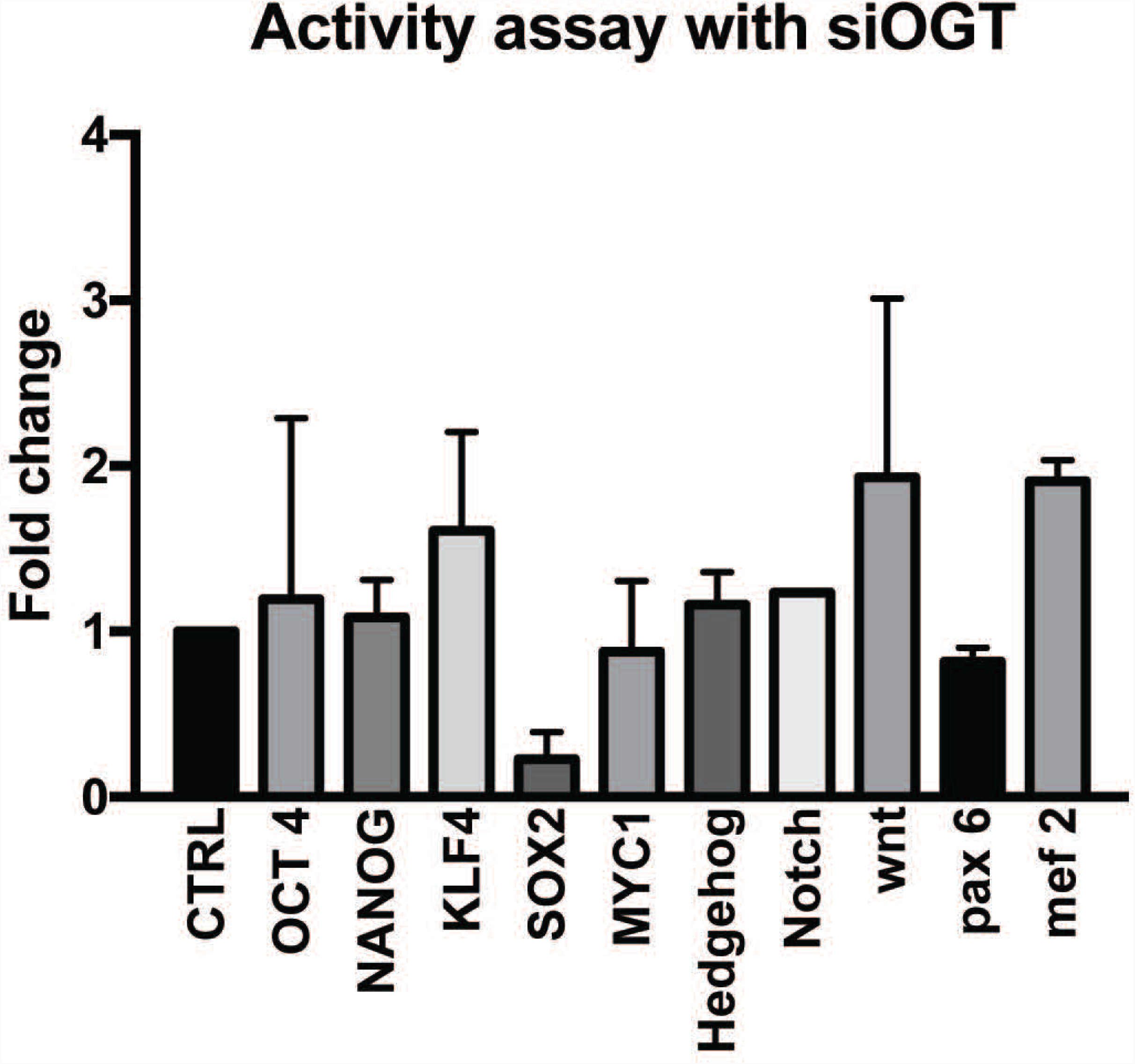
Luciferase reporter based transcription activity assay for stemness related transcription factors show that Sox2 activity was decreased upon inhibition of OGT by siRNA (A).

**Supplementary Figure 4.**
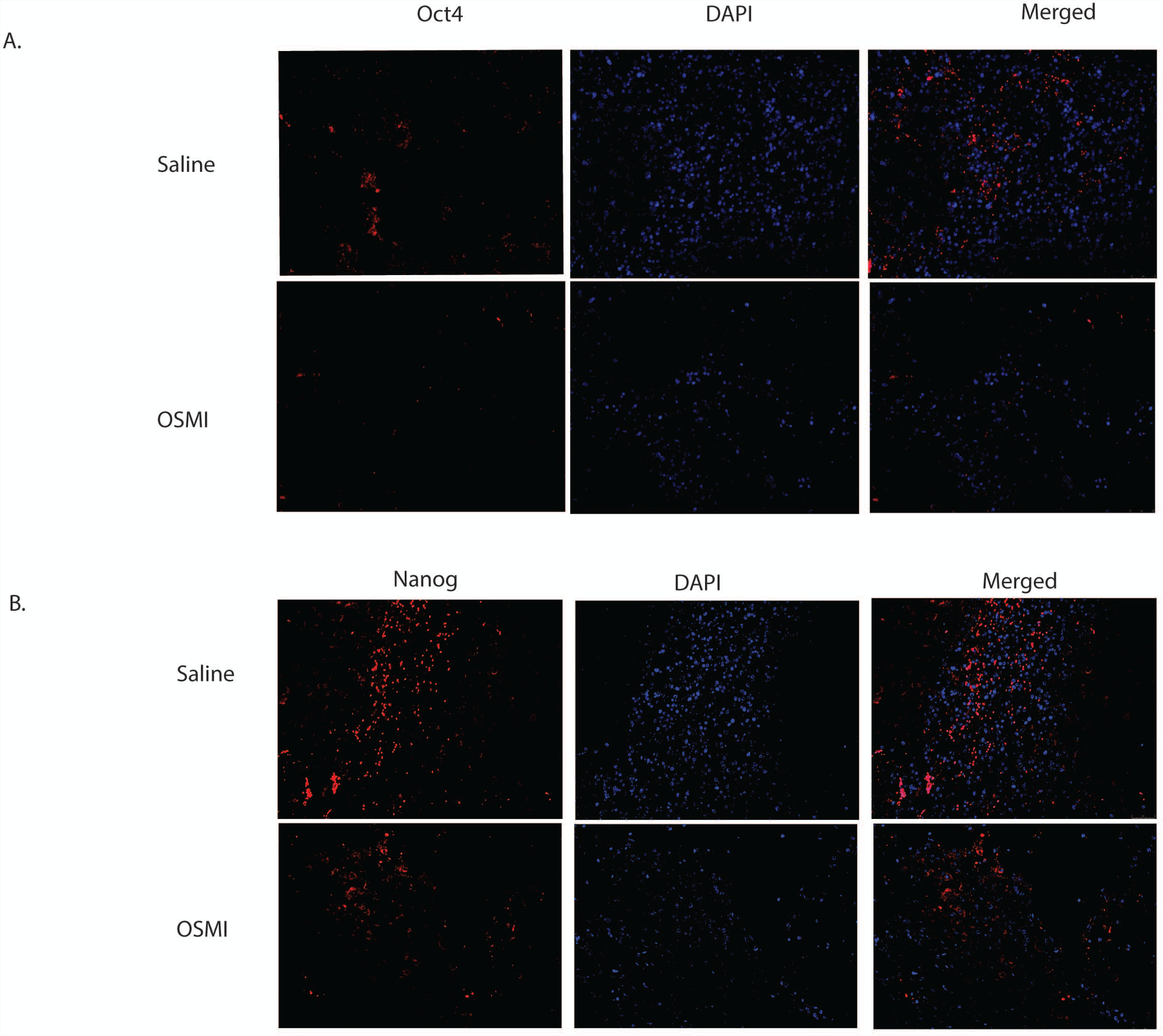
Treatment with OSMI decreases OCT4 expression (A) and NANOG expression (B) in the tumors compared to control.

